# Competitive ecosystems are robustly stabilized by structured environments

**DOI:** 10.1101/2020.03.09.983395

**Authors:** Tristan Ursell

## Abstract

Natural environments, like soils or the mammalian gut, frequently contain microbial consortia competing within a niche, wherein many species contain genetic mechanisms of interspecies competition. Recent computational work suggests that physical structures in the environment can stabilize competition between species that would otherwise be subject to competitive exclusion under isotropic conditions. Here we employ Lotka-Volterra models to show that physical structure stabilizes large competitive ecological networks, even with significant differences in the strength of competitive interactions between species. We show that for stable communities the length-scale of physical structure inversely correlates with the width of the distribution of competitive fitness, such that physical environments with finer structure can sustain a broader spectrum of interspecific competition. These results highlight the generic stabilizing effects of physical structure on microbial communities and lay groundwork for engineering structures that stabilize and/or select for diverse communities of ecological, medical, or industrial utility.

**AUTHOR SUMMARY:** Natural environments often have many species competing for the same resources and frequently one species will out-compete others. This poses the fundamental question of how a diverse array of species can coexist in a resource limited environment. Among other mechanisms, previous studies examined how interactions between species – like cooperation or predation – could lead to stable biodiversity. In this work we looked at this question from a different angle: we used computational models to examine the role that the environment itself might play in stabilizing competing species. We modeled how species arrange themselves in space when the environment contains objects that alter the interfaces along which competing species meet. We found that these ‘structured’ environments stabilize species coexistence, across a range of density of those objects and in a way that was robust to differing strengths of interspecies competition. Thus, in addition to biological factors, our work presents a generic mechanism by which the environment itself can influence ecological outcomes and biodiversity.

## INTRODUCTION

Natural environments from scales of microbes^1–4^ to large ecosystems^5–8^ are replete with communities whose constituent species stably coexist at similar trophic levels, despite apparent competition for space and resources. Generically, in spatially limited ecosystems species grow until resources and/or interactions with other species (e.g. competition or predation) limit their populations, notably not necessarily at a constant level through time^9–11^. In some cases^12^, the same set of species may exhibit qualitatively distinct relationships in a way that depends on available resources, with corresponding maintenance or loss of diversity. Species diversity and ecosystem stability have a complicated relationship^13,14^, and qualitatively different theories have been developed to explain variations in species diversity and abundance in resource-limited natural environments^15,16^. At one extreme, the principle of competitive exclusion asserts that if more than one species is competing within a niche, variations in species reproduction rates resulting from adaptation to the niche will necessarily lead one species to dominate within that niche to the exclusion of all other species, potentially driving inferior competitors into other niches^17–21^. Thus, competition for resources within a niche would push ecosystems toward lower species diversity^22^. At the other extreme are so-called ‘neutral theories’ which offer the null-hypothesis that organisms coexisting at similar trophic levels are – *per capita* – reproducing, consuming, and migrating at similar rates, and hence maintenance of biodiversity is tantamount to a high-dimensional random-walk through abundance space^23–25^. Such models often require connections to an external meta-community to maintain long-term stability^26^, lest random fluctuations will eventually drive finite systems toward lower diversity^27,28^. Many other mechanisms (which we cannot do justice to here) have also been proposed for maintenance of diversity in competitive ecosystems, including by not limited to: stochasticity and priority effects^29,30^; environmental variability^31^; models that encode specific relationships between species to maintain diversity^32^ (including the classic rock-paper-scissors spatial game^11^, cross-feeding^33–37^, metabolic trade-offs^38–40^, or cross-protection^41^); varied interaction models^42^; higher-order interactions – beyond pairwise – that stabilize diversity^43–46^; and systems where evolution and ecological competition happen simultaneously^47,48^. We do not take issue with any of these models / mechanisms, indeed, all of them are likely relevant and useful within certain contexts. Rather, this work uses computational modeling to argue that physical structure within an environment is a generic and robust mechanism for maintaining biodiversity in competitive ecosystems, across differences in competitive parameters, length-scales of physical structure, and for an arbitrary number of distinct species given lower-bound requirements on available space.

Microbial ecosystems present a particularly attractive test-bed for these ecological ideas. From a practical point of view, they are small and fast growing, relatively easy to genetically manipulate, and can be grown in controlled and customizable synthetic environments^35,49–52^, such as microfluidics^53,54^. Conceptually, characterizing the forces and principles that establish and maintain microbial biodiversity is of significant interest in health-relevant settings like the human gut^55–57^ and in the myriad contexts where soil microbiota impact natural or agricultural ecosystems^1,58^. These contexts motivate the model herein discussed, which can be conceptualized as a multi-species microbial ecosystem adherent to reasonable simplifications that make computations tractable. We used a spatial Lotka-Volterra model and assumed that all pairwise interspecies interactions were competitive. We focused on the role that physical structure of the environment plays in long-term dynamics of such ecosystems. Within the context of this model, our results were clear – steric structures distributed throughout the environment foster biodiversity in a way that depends less on the specific arrangement of those structures and more on the length scale of separation between the structures^59^. This structural stabilization was robust even when the ecosystem contained significant asymmetries in the competitive interactions between species, and the degree of stabilization positively correlated with decreasing structural scale. Finally, we provide evidence that the stabilizing effects of steric structure extend to an arbitrary number of species in competition with each other, as long as there is enough space.

## Results

### Competition and Structural Model

We modeled interspecies interactions using the canonical spatial Lotka-Volterra (LV) framework, with simplifying assumptions motivated by attributes of microbial ecosystems. For all species, we assumed that the carrying capacity per unit area of the environment is the same, that the basal growth rate *r* is the same, and that their migration can be described by random walks with the same diffusion coefficient *D*. Using those assumptions, the system of partial differential equations (PDEs) that describe an *N*-species LV model can be non-dimensionalized and, without loss of generality, written as

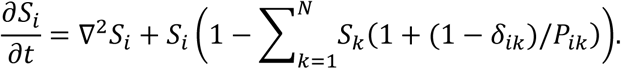

Each focal species *S*_*i*_ has a local concentration from zero to one in units of the carrying capacity, time is measured in units of inverse growth rate *r*^-1^ and the natural length scale 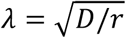 is proportional to the root mean squared distance an organism will move over a single doubling time. Self-interactions and simple competition for space are accounted for by the constrained carrying capacity and the corresponding sum over *S*_*k*_, and thus the diagonal elements *P*_*ii*_ are removed by the Kronecker delta, *δ*_*ik*_. Pairwise interactions between species are described by the off-diagonal matrix elements *P*_*ik*_ which are the concentrations of species *S*_*k*_ above which *S*_*k*_ actively reduces the concentration of *S*_*i*_. Neglecting intrinsic permutation symmetries, there are *N*(*N* − 1) pairwise interaction parameters for each *in silico* ecosystem, which can be thought of as a directed graph whose edges connect each pair of species. We focused on the class of ecological graphs that correspond to all species in competition with all other species, termed ‘all-to-all’ (ATA) competition, which corresponds to all off-diagonal elements *P*_*ik*_ > 0. This work uses computationally tractable values of *N* to support general claims about the dynamics of ATA ecosystems in structured environments, albeit such computations do not constitute a rigorous proof.

This model is appropriate for describing diffusively-mediated locally competitive interactions; examples of such ecosystems include situations where multiple species compete for the same pool of resources and actively reduce competitor abundances through (e.g.) Type VI secretion system mediated killing^60,61^, secretion of toxins ^62,63^ or antibiotic antagonism^64,65^. Analysis of bacterial genomes indicates that (at least) a quarter of all sequenced species contain loci encoding for mechanisms of active competition toward other species^66^ (though not necessarily for the purposes of consuming them as prey). Additional PDEs would be required to describe highly motile cells, chemotaxis in exogenous chemical gradients, or the production, potency, transport and decay of rapidly diffusing secreted toxins. This system of PDEs (which is not new^67,68^) represents a baseline set of assumptions and corresponding phenomena from which to build more complex models^69^ of structured population dynamics.

The *O(N*^2^) dimensional parameter space is too large to exhaustively sample for large *N*, and thus we employed statistical sampling techniques. We sampled a uniform random distribution of values for the off-diagonal elements *P*_*ik*_ whose mean was ⟨*P*⟩ = 0.25 and whose width was specified by the parameter Δ*P*. The value ⟨*P*⟩ = 0.25 indicates that on average when the local concentration of a given species reaches one-quarter of the carrying capacity, active reduction of neighboring competitors will occur. For each value of Δ*P* and structural parameters, we performed 30 to 50 simulations each with a unique random sampling of the interaction parameters *P*_*ik*_. All simulations had initial conditions in which every grid position had a low (0.2%) probability of being populated by any one of the available species, such that each species could grow and claim territory before competing. The data herein presented uses an 8-species system (56 interaction parameters), though in the last section we examine larger values of *N.*

Our *in silico* environments were square 2D planes with steric pillars distributed in the simulation space. Both the pillars and the bounding box were modeled with reflecting boundary conditions, thus, like a grain in soil or tissue in a gut, these pillars do not allow free transport through them, nor microbes to occupy them. Interspecies boundaries within the simulation area are primarily impacted by steric spacing^70^, and thus for simplicity the radii of the pillars were held fixed at *R* = 3 in dimensionless units for all simulations. We varied the mean distance between steric objects relative to pillar radius (Δ*x*/*R*), which we refer to as the ‘structural scale’, and we varied the degree of disorder in the arrangement of those steric objects. Disorder was introduced by translating each pillar in a random direction by an amount drawn from a uniform random distribution whose width is reported relative to the structural scale. Thus, disorder is characterized by a continuous dimensionless variable *δ*, which when equal to zero means the pillars are arranged in an ordered triangular lattice, and as *δ* increases the pillars approach a random arrangement in the simulation space (including the possibility of overlap) (see SI Fig. 1).

**Figure 1:**
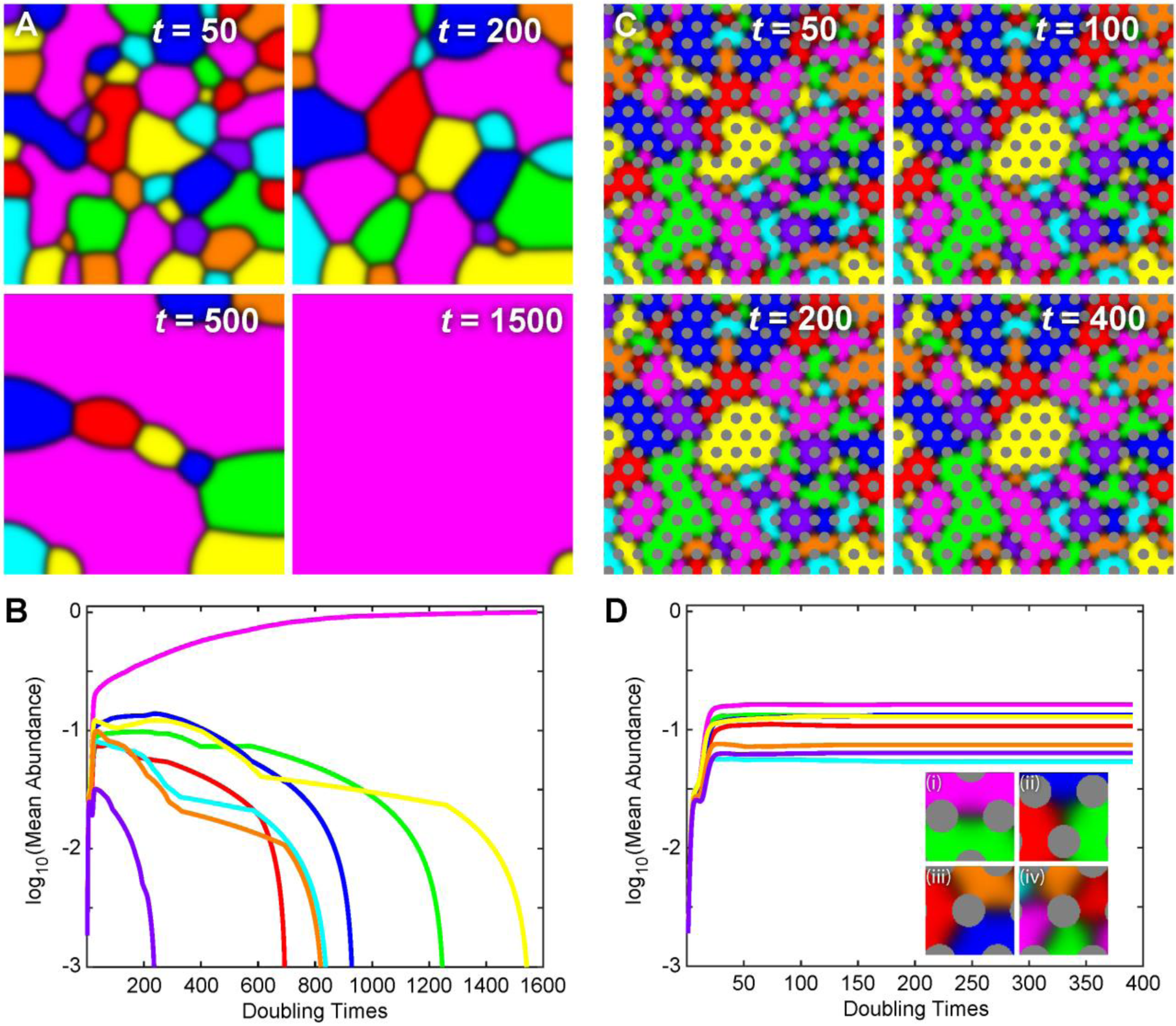
Steric structures stabilize an *in silico* multi-species ecosystem. Shown here are two 8-species *in silico* ecosystems in which all species are actively competing with all other species and the competition parameters are equal for all pairwise interactions. **(A & B)** Simulation of competition in an isotropic environment. If initial species representation is statistically equal, each species has equal probability of dominating the environment in the long-time limit. However, in any single simulation the dynamics of interspecies boundaries lead to a single dominant competitor in the long-time limit. **(C & D)** When species compete in an environment with ordered steric structures (*R* = 3, Δ*x/R* = 3.5, and *δ* = 0), interspecies boundaries that are mobile under isotropic conditions quickly ‘pin’ between steric objects and arrest the genetic-phase coarsening that leads to a single dominant competitor, thereby robustly producing stable representation of all species. Steric structures also permit situations where 3 or more species form a ‘junction’ around a steric object (Diii and Div). In both simulations *L* = 150, ⟨*P*⟩ = 0.25, and Δ*P* = 0.

Finally, it is worth noting limitations and simplifications inherent in this modeling framework. These systems of PDEs are deterministic, that is, with the same interaction parameters, simulation size and initial conditions they produce the exact same dynamics. Stochasticity enters our simulation framework through the random initial conditions, disorder, and random samplings of the interaction parameters. A number of excellent studies have examined low*-N* systems using fully stochastic dynamics^71–75^, revealing quantitative differences between deterministic and stochastic models, as well as qualitative differences over long time scales where stochastic fluctuations can drive a system into new parts of phase space, for example, into extinction cascades^76,77^ or mobility-dependent biodiversity^71,73,78^. Details of our computational setup are discussed in the Methods section and all of our code is available upon request.

### Environmental structure stabilizes all-to-all competition

We compared the spatial population dynamics of an 8-species system with and without the inclusion of spatial structure. In both cases, each species engaged in active (population reducing) competition with every other species, for a total of 56 pairwise interactions each characterized by the value of an off-diagonal matrix element. In Fig. 1 we show the simplest version of this comparison, where the strength of the competitive interaction is equal between all pairs of species (i.e. all off-diagonal elements have the same value, Δ*P* = 0). When a system lacks steric structure, and hence is spatially isotropic, a single dominant competitor will emerge to the exclusion of all other species^79^ (Fig. 1 A/B). Spatial population dynamics are determined by an interplay between the curvature of the interface between any two species and the relative values of the competition parameters for the species that meet at that interface^70^. If competition at a particular two-species interface is balanced (i.e. *P*_*ik*_ = *P*_*ki*_) then interfacial curvature is the only determinant of interface movement; straight interfaces do not translate and curved interfaces translate in the direction that straightens them. However, if the interaction at a two-species interface is unbalanced (i.e. *P*_*ik*_ ≠ *P*_*ki*_) then there is a critical interface curvature below which the stronger competitor will invade the space of a weaker competitor, ultimately to the exclusion of the weaker competitor. Excepting literal edge cases, wherein boundaries between species contact multiple parallel edges of the simulation space, the dominance of a single competitor in isotropic space is robust to changes in the size of the simulation space, the number of species and the values of interaction parameters, given enough time.

In contrast, the inclusion of physical structure leads to long term, stable representation of multiple (and often all) species across a range of structural scales and interaction parameters. In Fig. 1 C/D we show the evolution of balanced competitive interactions between 8 species with the same initial conditions and interaction parameters as Fig. 1 A/B, but in the presence of a triangular lattice of steric pillars. In this system, the abundances of all 8 species rapidly equilibrated leading to stable coexistence. In this stable state, each species established spatial domains whose boundaries were primarily composed of two-species interfaces governed by the same rules of interfacial curvature and competitive parameters discussed above (Fig. 1Di). The steric pillars also stabilized a number of distinct ‘junctions’ between species, including free three-way junctions in open space (Fig. 1Dii), three-way junctions centered on a pillar (Fig. 1Diii), and four-way junctions centered on a pillar (Fig. 1Div). Isotropic systems can transiently support two-species interfaces and free three-way junctions (Fig. 1A), but the three- and four-way junctions centered on a pillar can only exist in systems with steric objects. Junctions centered on pillars can support more than four species if pillars are large in comparison to the length scale (thickness) of the interface, though these did not occur in our simulations with random initial conditions – we only observed these higher-species junctions under contrived conditions (see SI Fig.2).

### Stabilization is robust to fluctuations in structure and competition asymmetries

Next we held the degree of structural disorder fixed at zero (*δ* = 0) and explored how changes in the structural scale affected the number of stably coexisting species. Along one dimension, we held ⟨*P*⟩ = 0.25 and varied the interaction parameter Δ*P* subject to the constraint that Δ*P*/2 < ⟨*P*⟩, which ensured that all *in silico* ecosystems remained in the ATA graph class. Along a second parametric dimension, we varied the structural scale while holding pillar radii fixed. In Fig. 2 we measured the mean number of species at equilibrium as a function of both the width of the interaction-parameter distribution and the structural scale, with 30 replicates per parameter set for a total ∼11,000 simulations. Consistent with previous work on two-species systems^70^, the average number of surviving species declined sharply both as competition asymmetry increased (i.e. as Δ*P*/⟨*P*⟩ increased) and as the structural scale increased. Conversely, systems with smaller structural scales and/or smaller competitive asymmetries robustly retained all eight species in the long-time limit. The structural scale sets the maximum interface curvature that can exist in an ordered environment, and hence limits the values of Δ*P*/⟨*P*⟩ for which all species can survive, that relationship is given by

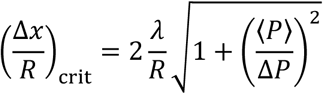

derived by setting the maximum geometric curvature equal to the curvature due to competitive asymmetry (see SI of ^70^). This relationship approximates the boundary between the regime of stable coexistence of all species and reduced species coexistence, as shown overlaid on Fig. 2.

**Figure 2:**
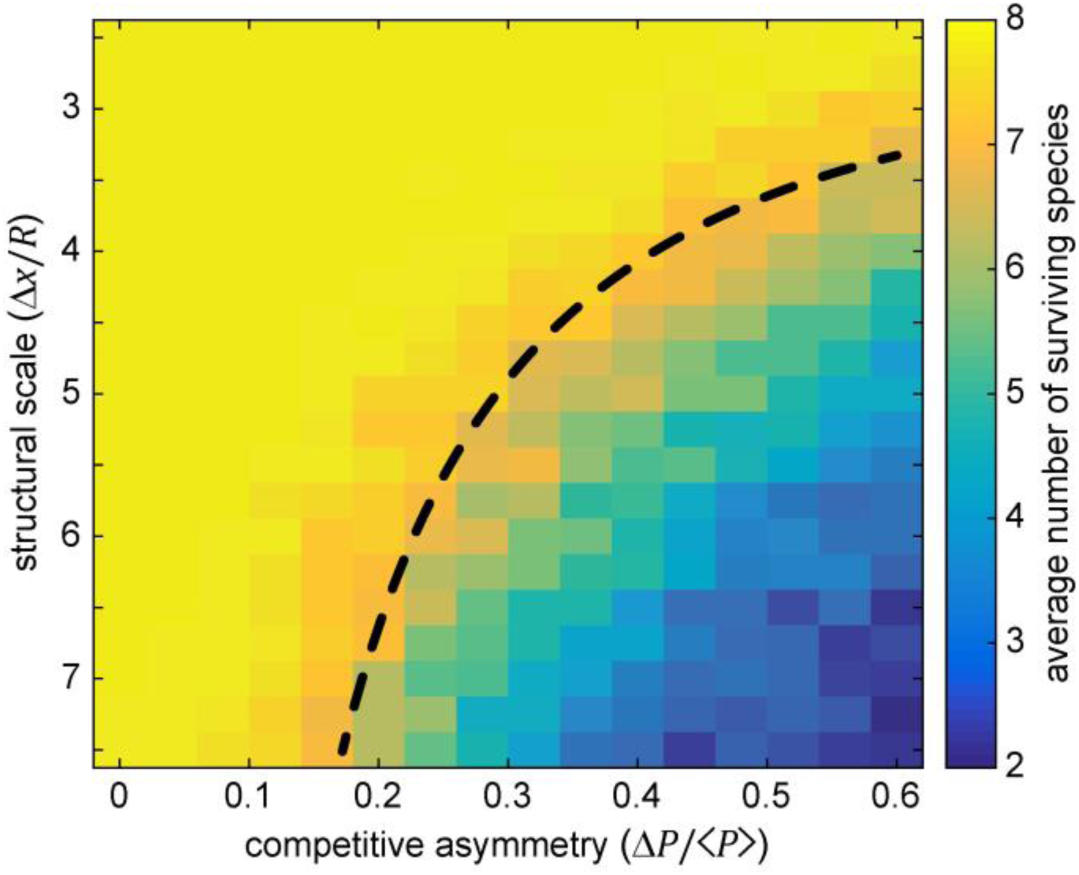
The number of species an environment can stably support depends on the degree of competition asymmetry and the structural scale. Simulations were performed across a range of competitive asymmetries characterized by the dimensionless parameter Δ*P*/⟨*P*⟩ and across a range of structural scales characterized by the dimensionless parameter Δ*x /R*. Each pixel corresponds to the mean of 30 simulations each with a unique random sampling of initial conditions (as described in Methods), a unique random sampling of the interaction matrix elements using Δ*P* and ⟨*P*⟩, and a fixed structural scale. Structural scale and competitive asymmetry were both potent modulators of species coexistence, with smaller structural scale and lower competitive asymmetries leading to stable representation of all species (yellow region). The black dashed line is a zero-fit parameter model relating the competitive asymmetry to the maximum curvature possible for a given structural scale, showing that relatively simple geometric considerations capture the onset of species loss. In all simulations *L =* 150, ⟨*P*⟩ = 0.25, *R =* 3, and *δ* = 0.

While these results are supportive of the stabilizing effect of steric structure on long-term species coexistence, rarely do natural environments contain highly ordered (*δ* = 0) steric structures, thus we explored how disorder affects species abundance at a fixed structural scale. First, we simulated the simpler case where all eight species had balanced competitive interactions (like Fig. 1) and examined the population dynamics in the presence of disordered steric structures (*δ* = 1). Like the ordered case, disordered systems with balanced competitive interactions displayed stable representation of all eight species (Fig. 3A). We then compared the probability distribution for the number of coexisting species at equilibrium across four conditions (1,000 simulations for each): with and without competitive asymmetry, and with and without structural disorder, as shown in Fig. 3B. When Δ*P*/⟨*P*⟩ = 0 the number of species remained at the maximum value across all 1,000 simulations whether or not the steric structures were disordered. When competitive asymmetry was introduced the probability distribution for the number of stably coexisting species expanded across all possible values (1 to 8) and peaked between 6 and 7 species, regardless of whether the system was ordered or disordered. Those distributions were quantitatively similar, indicating that disorder was not a strong determinant of stable species coexistence.

**Figure 3:**
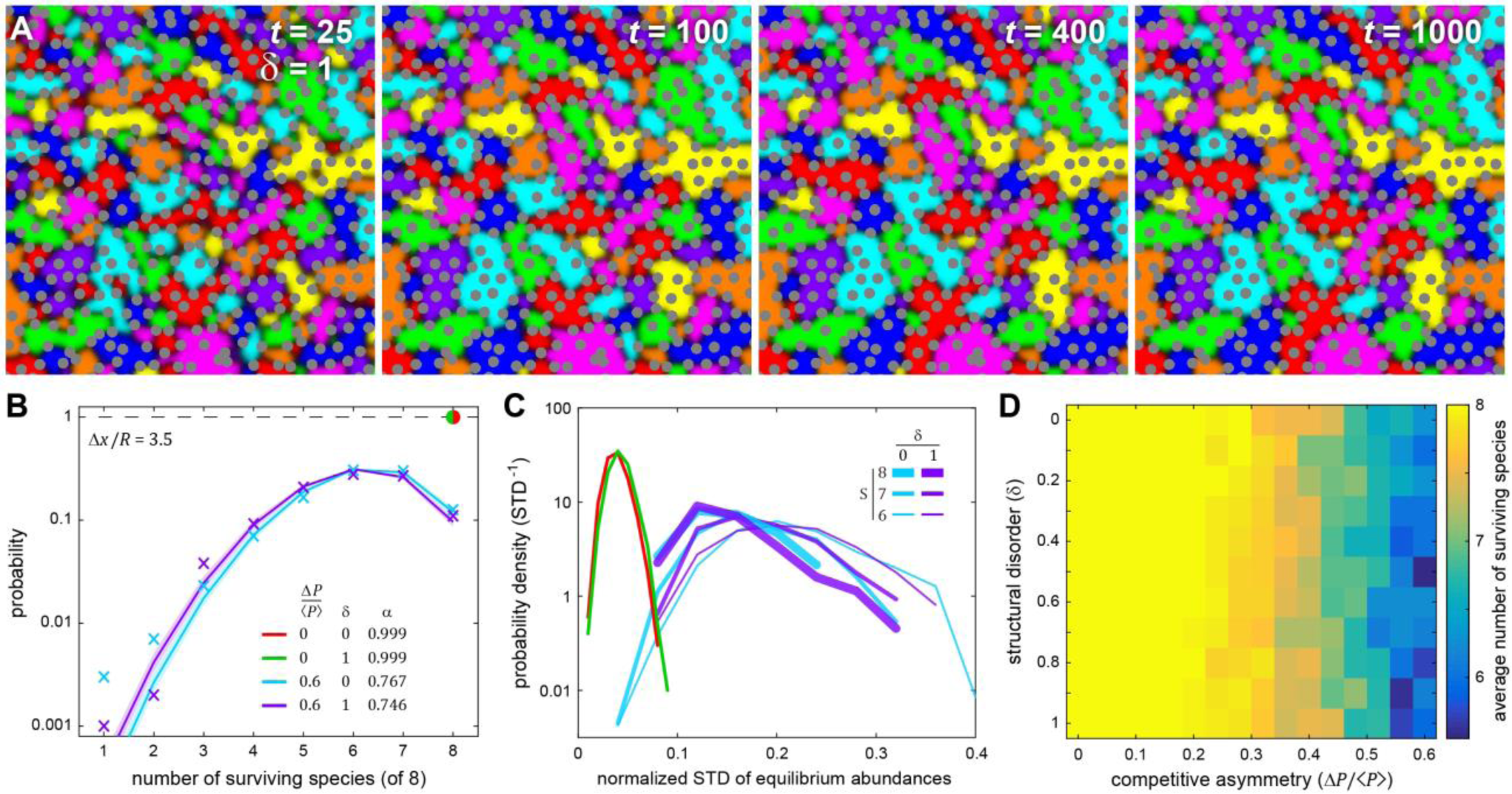
Species coexistence is robust in the presence of structural disorder. **(A)** Similar to Fig. 1C, these panes show the time evolution of an 8-species system where steric pillars were randomly perturbed from a perfect triangular lattice (*δ* = 1). From random initial conditions the system rapidly equilibrated to a stable configuration that supported all 8 species. **(B)** Simulations were performed to measure the probability distribution for the number of surviving species under four conditions (1,000 each): high and low competitive asymmetry and high and low structural disorder. The structural scale was held fixed. Without competitive asymmetry all species survived in all simulations, with or without disorder (red/green dot). With high competitive asymmetry, the probability distributions spread across all possible numbers of species with relatively little distinction between ordered and disordered systems (colored X’s). Those distributions were well-described by a modified binomial distribution (solid lines) with an ensemble average single-species survival probability of *α* ∼ 0.75. The distributions exhibit larger variation where there are less data (i.e. lower probabilities, from one to three surviving species). **(C)** For each simulation, the normalized standard deviation (NSD) in equilibrium abundances was measured, and those values are shown here as a histogram for each of the four conditions. In the absence of competitive asymmetry (green, red), the mean NSD was low (∼0.05 on maximum scale of 1). When competitive asymmetry was high, we examined the NSD for all simulations that had 6, 7, or 8 surviving species (cyan, purple). All of those distributions were significantly wider and had significantly higher mean NSD’s, meaning that competitive asymmetry increases the variation in species abundance at equilibrium, regardless of how many species stably coexist. **(D)** We compared the average number of surviving species across a range of competitive asymmetry and disorder (0 ≤ *δ* ≤ 1). Those distributions showed little variation with disorder and, as a function of competitive asymmetry, had a mean pairwise correlation of 0.97 (see SI Fig. 4). Thus our data suggest that structural scale has a stronger effect than disorder (at least over this range). In all simulations *L =* 150, ⟨*P*⟩ = 0.25, and *R =* 3.

We examined the survival probability distributions and applied the simplest possible rule that emerges from the statistical ensemble of initial conditions, competitive asymmetries, and disorder. For a given set of conditions we assumed that there is some probability *α* that a species randomly selected from the full ensemble will survive in the long-term. This corresponds to a binomial distribution whose normalization is modified to account for the fact that there is no chance that all *N* species will die

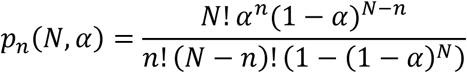

where *N* is the maximum (initial) number of species and *p*_*n*_ is the probability of 1 ≤ *n* ≤ *N* species coexisting at equilibrium. We used maximum-likelihood estimation to fit this model to the survival distributions and thus determined the value 0 < *α* < 1 that corresponds to the ensemble average probability that any single species survives in equilibrium given the number of species *N*, simulation size *L*, structural parameters Δ*x* and *R*, the disorder *δ*, and sampling parameters ⟨*P*⟩ and Δ*P*; the exact functional dependence between those parameters and *α* is not yet clear. This model captures the bulk of the survival distributions and mis-estimates the occurrence of rare events compared to the raw data. The fit values demonstrate that smaller systems and systems with higher competitive asymmetries both have lower per-capita survival probabilities (*α*). For instance, examining systems with higher competitive asymmetry, we found that larger systems (*L* = 150) with or without disorder had quantitatively similar survival probabilities – 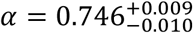 and 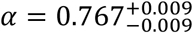 respectively – whereas under the same conditions a smaller system (*L* = 75) had survival probabilities of 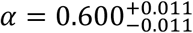 and 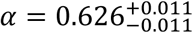, respectively (See SI Fig. 3).

We used those same 4,000 simulations to assess the variability in the amount of area that each species occupied at equilibrium divided by the total simulation area, giving a dimensionless quantity bounded between 0 and 1 that characterizes variability in species abundance. For each set of parameters, we measured the standard deviation in abundance across all species and all replicates to generate a probability distribution for the degree of variation – values closer to zero indicate that all species have roughly the same abundance. In the case of balanced competitive interactions, the probability distributions for abundance variations were nearly identical for the ordered and disordered systems, and the mean value of the variation was low (∼0.05), meaning that if competitive interactions are balanced all species are have roughly the same abundance. However, introducing moderate competitive asymmetry meant that some species intrude into the territory of other species, leading both to lower species diversity and to larger variations in species abundance. As such, we report the distribution of abundance variations for all simulations that had equilibrium species numbers of *S* = 6, 7 and 8 (Fig. 3 C). Again, disorder had little effect on those probability distributions. Systems that experienced extinctions of zero (*S* = 8), one (*S* = 7) or two (*S* = 6) species had a higher mean variation (by a factor of 3 to 5) and wider distribution of variations as compared to balanced competition. Additionally, we performed a wider sampling of the degree of disorder and the competitive asymmetry, and found that the average number of stably surviving species showed little dependence on the degree of structural disorder (Fig. 3D). When we correlated the mean number of surviving species as a function of the competitive asymmetry across all values of *δ*, the average correlation coefficient was 0.97 (see SI Fig. 4), meaning that the relationship between average number of surviving species and competitive asymmetry showed little dependence on disorder in the range 0 ≤ *δ* ≤ 1.

**Figure 4:**
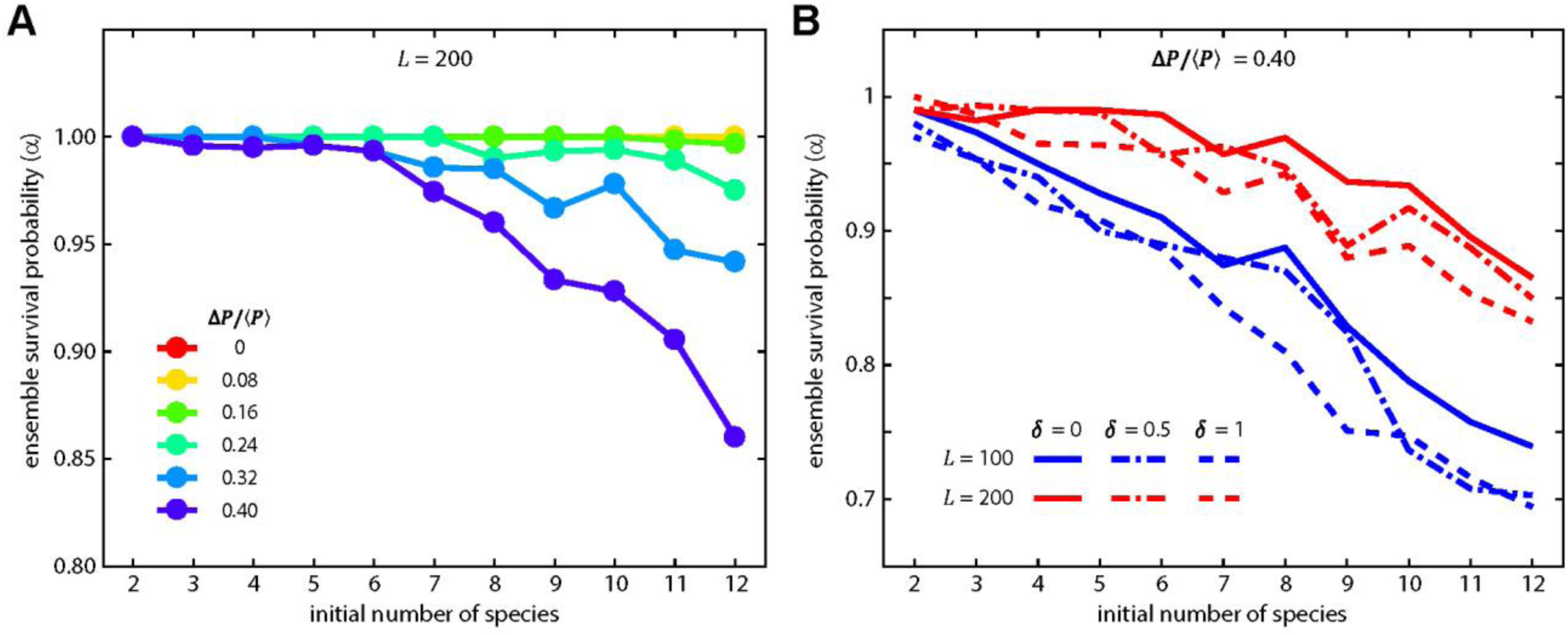
Structure stabilizes larger numbers of species, but increasing competitive asymmetry increases species loss. **(A)** Holding the structural scale fixed with no disorder, we measured the survival probability as a function of the initial number of species, between 2 and 12, across multiple values of competitive asymmetry. For lower values of competitive asymmetries, the final and initial numbers of species were essentially equal (*α* ∼ 1). Higher levels of competitive asymmetry resulted in increasing degrees of species loss as the number of species increased. This amplification of species loss is related, at least in part, to the same interplay between structural scale and maximum interface curvature that caused species loss in Fig. 2. **(B)** Comparing identical conditions between two different system sizes (*L =* 100 and *L =* 200), the effects of disorder are relatively small in comparison to the effects system size, with smaller system sizes (blue lines) showing a significant amplification of reduction in survival probability as compared to the larger system. In all simulations ⟨*P*⟩ = 0.25 and *R =* 3.

As a final characterization of spatial structure, we examined the density with which interspecies boundaries connect steric objects. Ignoring edge cases, every steric object has a set of Voronoi nearest neighbors, typically 5 to 7 in disordered systems and exactly 6 in a triangular lattice (see SI Fig. 5). Across our simulations, the vast majority (∼ 98%) of all interspecies boundaries connected pillars that were Voronoi nearest neighbors, which is expected given geometric constraints. The number of those connections per unit area relative to the total number of possible Voronoi connections per unit area is a dimensionless measure of the amount of competition in a physically structured system – below we refer to this as the ‘connection density’, whose values lie between 0 and 1. When examined through that lens, disorder, at least for balanced interactions, had a significant effect on the distribution of connection densities across the ensemble of simulations, with ordered systems exhibiting higher connection densities (see SI Fig. 5). However, when examined under moderate levels of competitive asymmetry connection density significantly decreased (consistent with abundance variation increasing) and the difference between ordered and disordered systems again became small. These data suggest that structured systems with higher levels of competitive asymmetry, somewhat counterintuitively, have lower levels of competition as measured by the density of competitive interfaces in the system, because boundaries between mismatched competitors are less stable.

### Structure stabilizes larger numbers of species with system-size dependence

All of the simulations discussed up to this point were performed within the same size grid *L* = 150 (with the exception of SI Fig. 3, *L* = 75). Under any set of parametric conditions, the absolute minimum domain size for a given species is set by the area of a triangle formed by three pillars that are all mutual Voronoi neighbors (so-called ‘Delaunay triangles’^80^), thus *any* system that is not large enough to contain domains of at least that size for each of *N* unique species cannot support all *N* species – this establishes a minimum system size for a particular number of species that scales as *N*(Δ*x*)^2^. However, disordered systems have an additional system-size dependence – all else being equal, as the system size grows the probability distribution for the equilibrium number of species (e.g. Fig. 3B) shifts toward the maximum number of species (i.e. *α* → 1). The mechanism behind this shift is that as the system increases in size, there are more opportunities for disordered steric objects to create a zone in which a weaker competitor is enclosed by an effectively smaller structural scale. We confirmed this by measuring the system size-independent distribution of local structural scale (SI Fig. 6). In an ordered system, the distribution of local structural scale is a delta-function centered on the lattice constant. As disorder increased we found that Voronoi zones emerged whose maximum convex edge-length was smaller than the lattice constant, meaning these were zones in which a species that would be too weak to compete in an ordered system, could potentially survive. The average number of these zones per unit area is scale-independent, thus increasing system size linearly increases the average number of those zones, and thus the per-capita survival probability *α* increases (e.g. compare Figs. 3B and SI Fig. 3), as does the survival probability of weaker competitors.

Finally, in an ordered system we explored the effects of system size and competitive asymmetry as a function of the initial number of species, across the range 2 ≤ *N* ≤ 12. First, we measured the ensemble survival probability (*α*) as a function of both competitive asymmetry and the initial number of species, with 50 replicates for each parameter set (Fig. 4A). For low competitive asymmetry across all values of *N*, the ensemble survival probability was approximately 1, meaning all species survived, and hence the initial and final species numbers were equal. However, similar to Fig. 2, as competitive asymmetry increased species loss increased (*α* decreased), and the decrease in *α* was more dramatic the larger the initial number of species. One potential mechanism behind this species-number dependent change in *α* is that larger values of *N* offer a wider sampling of the matrix elements *P*_*ik*_, and thus increase the likelihood that a single competitor dominates over many other weaker species and/or that ‘ultra weak’ competitors emerge.

Another potential link between species-number and decreasing *α* was system-size. To test this, we ran simulations across the same range of species number at the highest value of competitive asymmetry for two different system sizes (*L* = 100 and *L* = 200). For both system sizes, survival probability dropped as species number increased (Fig. 4B), but the smaller system experienced a significantly faster drop in survival probability with species number. This indicates that decreasing system size accounts for part, but not all, of the decrease in survival probability. We also examined the role of disorder in this context; consistent with our previous results disorder had a relatively minor effect as determined by 95% confidence intervals of MLE for the values of *α* (see SI Fig. 7 for confidence intervals). Together, these results add support to the finding that disorder at a fixed structural scale is not a strong determinant of stable biodiversity, but that system size and the number of initially competing species each modulate survival probability and hence equilibrium biodiversity.

## DISCUSSION

In this work we used simulations to provide evidence that – within a range of structural scales and competitive parameters – the class of ecological graphs encompassed by all-to-all (ATA) competition is stable in structured environments. Other well-known ecological ‘games’, such as rock-paper-scissors (RPS) and its higher species-number analogs^73,81^, also reside within the ATA graph class. That is, values in the matrix *P* that produce stable intransitive (e.g. RPS) oscillations adhere to the same conditions as the ATA class, but are subject to additional constraints on their relative values. The RPS sub-class distinguishes itself by exhibiting two important features. First, unlike non-oscillatory systems that lie within ATA, oscillatory systems with spatial isotropy can exhibit stable representation of all species^43,71,82^, albeit with each species in constant spatial flux. Second, our previous work showed that in a symmetric game of RPS^70^, structure could have a destabilizing effect that ultimately led to extinction cascades ending with a single dominant species. Thus, while the vast majority of parameter combinations for *P*_*ik*_ likely yield systems that are stabilized by structure, RPS graphs present the possibility of being destabilized by structure. Given that virtually all natural environments present structural anisotropy, destabilization of RPS networks due to structure may contribute to answering why RPS networks are only rarely observed outside of the lab^21,83–85^.

Similarly, systems in which species cooperate or have competitive alliances – neither of which lie within the ATA graph class – can, by virtue of the specific localization of each species, be stabilized by structure. For instance, consider the simplest, non-ATA graph for which this can be true: *A* and *B* compete, *B* and *C* compete, and *A* predates *C* (see SI Fig. 8A). If in a structured environment species *C* is stably and spatially isolated from *A* by the arrangement of *B*, then all three species will stably coexist due to the effects of physical structure, even though an interspecies boundary between *A* and *C* is unstable regardless of structural parameters. The idea that specific spatial arrangements of species can be stable in a structured environment extends to other non-ATA graphs (e.g. SI Fig. 8B), and is consistent with established findings that spatially structured communities maintain biodiversity by localizing interactions among community members^86–88^. This also suggests the possibility that physical structure and positioning of species play a role in shaping their ecological and evolutionary relationships. Thus assessing the interplay between physical structure, graph structure, and ecological dynamics is a rich area of inquiry, one in which structure may play a qualitatively important role.

For simplicity and ease-of-display we explored 2D systems in this work, however many natural systems are three dimensional. This work does not allow us to directly comment on what will happen in 3D systems, however: (i) graph structure and its attendant parameters as encoded by the interaction matrix are independent of dimensionality, (ii) the relationship between interface curvature and competitive asymmetry that underlies many of the results herein described have a natural extension into three dimensions, where the mean curvature of the 2D interspecies boundary in 3D space plays the analogous role as 1D curvature of a line interface in 2D space, and (iii) the measures herein described (e.g. dimensionless disorder, structural scale, connection density, survival probabilities, etc.) have natural extensions into 3D, meaning that direct comparisons can be made between 2D and 3D systems. Similarly, there are illuminating comparisons to be made with other physical and biological systems, in particular the pinning phenomena that here slow or halt genetic coarsening play important roles in domain-wall stabilization in Ising-like systems due to pinning at random spatial impurities^89^, pinning-induced transitions of super-cooled liquids into glassy states^90^, arrest of lipid-bilayer domain coarsening in the presence of biopolymers that impose structure on the bilayer^91^, and have been shown to impact genotype-specific range expansion^92^. Likewise, other physical mechanisms, such as flow^93^, have been found to slow or halt coarsening in phase-separating systems, and still other work focuses on flow^94,95^ or chemical gradients^96^ in structuring communities. Thus structure is one of multitude physical mechanisms shaping communities in complex environments.

Finally, even within this 2D reductionist framework we wonder how robust are pinned competition interfaces to: (i) stochastic spatial fluctuations caused either by finite organism size or other forms of motility (besides diffusion), (ii) tunable interaction strengths, such as with competition sensing ^97,98^, or (iii) phenotypic differentiation ^99^? Whether the details of interactions matter^69^ or not^100^, the stabilizing effect of structure is fundamentally related to how interspecies boundaries – which appear in many extensions of the LV model – interact with the boundary conditions imposed by structure, and hence we anticipate that qualitatively, these results extend to a wider class of boundary-forming models (e.g. incorporating Allee effects^70^). Ultimately, an understanding of the interplay between ecological relationships, environmental structure, and other physical factors (like flow or chemical gradients) paves the way toward rational design of structured environments that tune the range of competitive asymmetries and/or stochastic fluctuations that an environment can stably support, and shift system dynamics and stability to favor particular species or interaction topologies.

## METHODS

In the 2D simulation space, each species was seeded by randomly choosing 0.2% of all valid pixels in the simulation box and setting the concentration of that species to 1/*N* of the carrying capacity, where *N* is the number of species being simulated; each species was represented by its own field matrix. Steric pillars were placed with the specified radius *R*, spacing Δ*x*, and degree of disorder *δ* within a simulation box whose side lengths were *L*. All reported length measures (*R, L*, Δ*x*) are in units of 1.29*λ*, with the computational pixel scale set so that 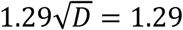 *λ* = 5. Microbial density that coincided with pillar locations was removed from the simulation. The bounding box and pillar edges were modeled as reflecting boundary conditions. At each simulated time step (Δ*t* = 0.01*t*, with *t* in inverse growth rate), populations diffused via a symmetric and conservative Gaussian convolution filter with standard deviation set by the diffusion coefficient, 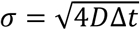. After the diffusion step, changes in population density (growth and death) were calculated using the equations given in the main text, and used to update the density of each species matrix according to a forward Euler scheme. In combination with the small dimensionless time-step and concentration-conserving convolution filter, hard upper and lower bounds (1 and 0 in units of carrying capacity, respectively) were enforced on each species field to ensure numerical stability of simulations; populations densities outside this range were set to 1 and 0, respectively. We monitored a subset of the simulations and found that simulations were sufficiently stable that those hard limits were never encountered. For each set of structural scale (Δ*x*/*R*) and competitive asymmetry (*P*_*ik*_) values, 30 to 50 independently initialized replicates were simulated for 5000 doubling times or until equilibrium was reached, as defined by spatially averaged change in all species matrices falling below 0.001 between time steps. Mean population abundances of the simulation were recorded at an interval of 100 Δ*t* for the duration of the simulation. Extinction was defined as the mean population density of any species dropping below a threshold value of ((2*R*)^2^ − *πR* ^2^)/4*A*, where *R* is the pillar radius and *A* the area of lattice points not obstructed by pillars. This non-zero threshold accounts for surviving populations ‘trapped’ between a pillar and the corner of the simulation box and therefore not in contact with the rest of the simulation; this threshold value lies well below the minimum set by the local Delaunay triangulation.

To create a controlled level of pillar-position disorder, each pillar center was displaced from a triangular lattice in a random direction by an amount drawn from a uniform random distribution whose width is characterized by the dimensionless parameter defined as *δ* = 2*w*/(Δ*x* − *R*), where *w* is the width of the uniform distribution. The competition parameters between all species were generated by choosing the mean strength of competition ⟨*P*⟩ and then sampling a uniform random distribution of width Δ*P* about that mean for each directed interaction (i.e. the interaction matrix is not symmetric), subject to the constraint that Δ*P*/2 < ⟨*P*⟩, which ensures that all random samplings remain in the ATA graph class.

### Model Fitting

Fitting to the modified binomial distribution was performed using maximum-likelihood estimation with all 1,000 simulations for each set of conditions; reported uncertainties for *α* are 95% confidence intervals.

### Data availability

Code files to run simulations and analyses are available as supplemental files that can be downloaded on the publisher website. Raw simulation output is available upon request.

## ACKNOWLEDGEMENTS

We thank Nick Lowery for helpful discussions and comments on the manuscript and Rob Yelle for assistance with high performance computing resources at the University of Oregon. Research reported in this publication was supported by the University of Oregon.

## Supplementary Information

**SI Figure 1.**
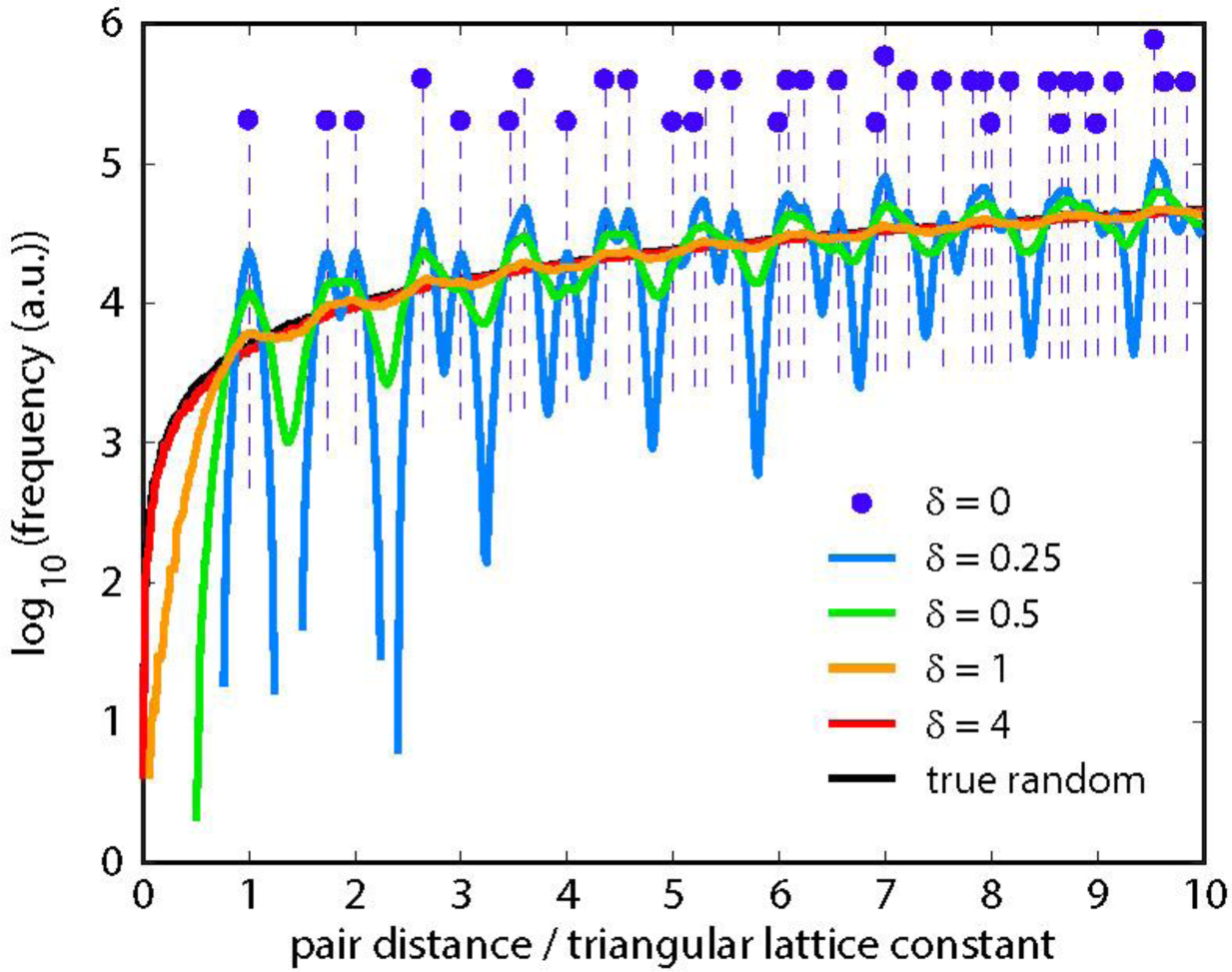
The pairwise distance distribution as a function of the disorder parameter *δ.* The distribution of separations between any two steric pillars does not depend on the size of the space in which those objects exist, assuming that the density of those objects is held fixed. For a perfect triangular lattice (*δ* = 0), that distribution is a series of delta-functions (purple dots). Adding structural disorder (*δ* > 0) makes the distributions continuous, with increasing degrees of disorder ultimately approaching the distribution expected for randomly placed objects at a fixed density (black line). As *δ* increases, the arrangement of steric pillars transitions smoothly from a triangular lattice to a random arrangement – in this work, we explored 0 ≤ *δ* ≤ 1.

**SI Figure 2.**
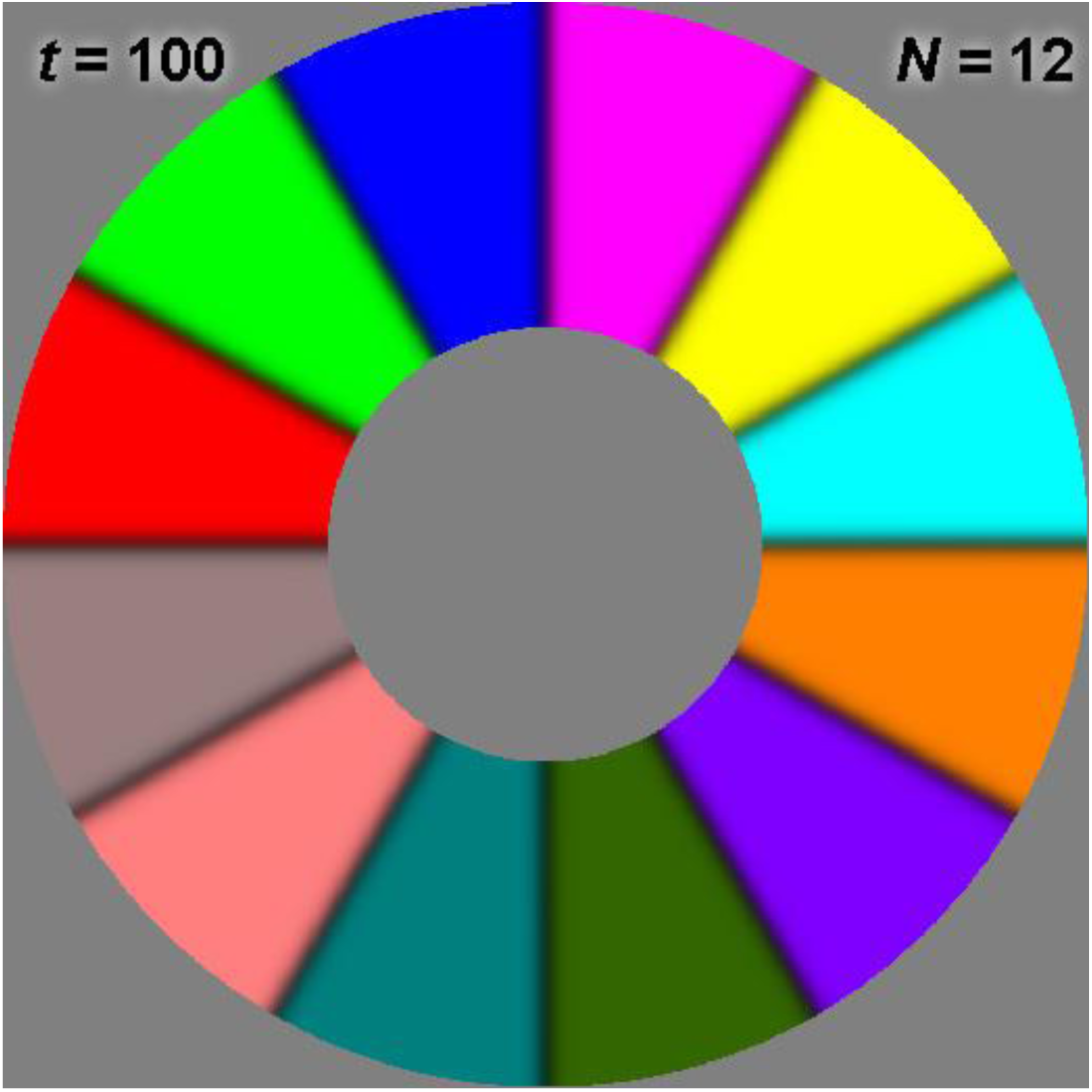
Here a simulation is contrived via initial conditions to have 12 species stably coexist, each making contact with the same steric object (gray circle in the center). The outer edge is also circular which permits stable coexistence of multiple species in a single open space. This figure demonstrates that, for a sufficiently large steric object as compared to the width of the interspecies transition zone, any number of species can ‘share’ a boundary with an object. However, in an environment with many steric objects in proximity, the local Voronoi neighborhood limits the maximum number of species that can exist around an object, usually to 3 or 4.

**SI Figure 3.**
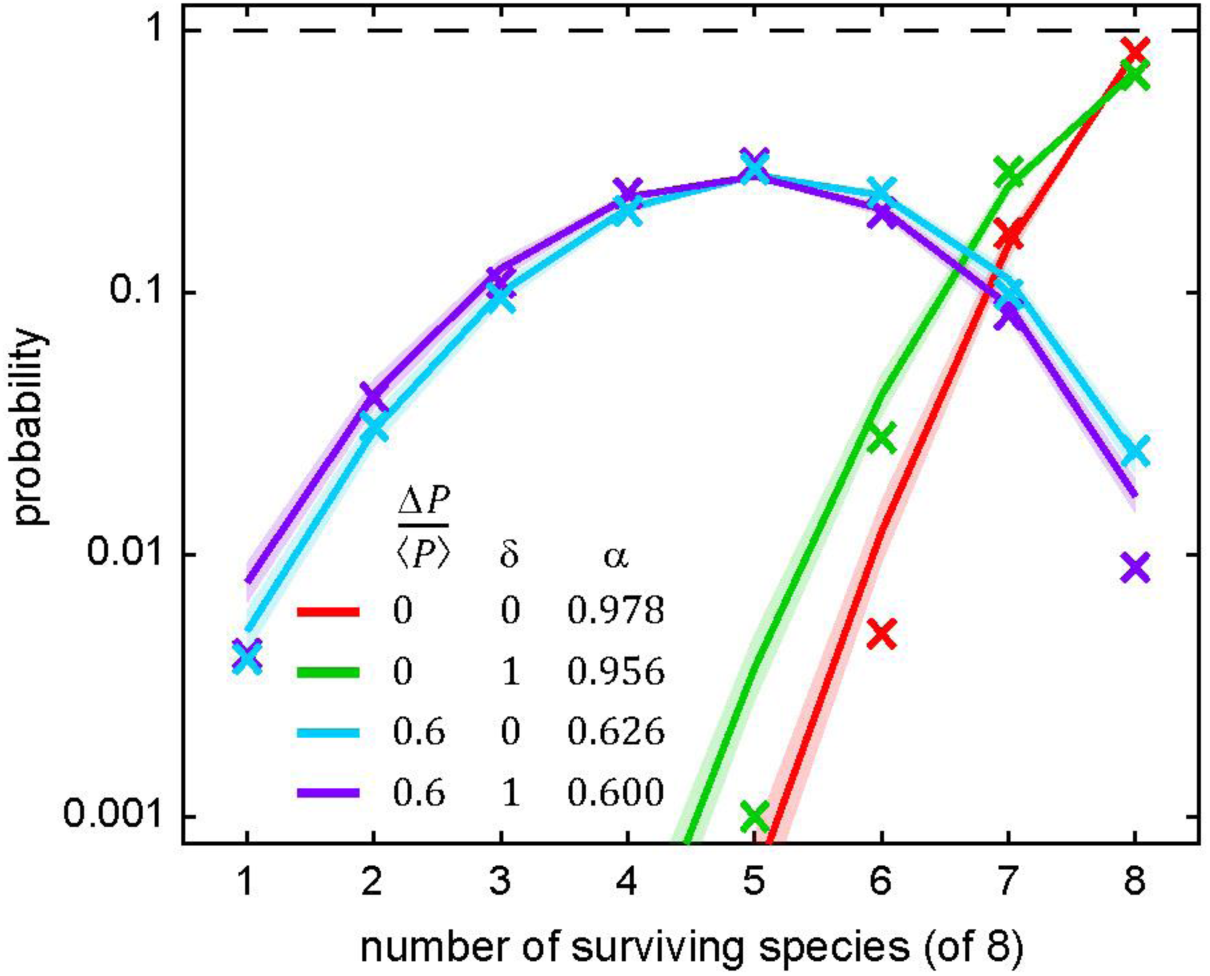
The type and format of data herein presented are the same as in Fig. 3B – the only difference is that here the simulation box is half the linear size (1/4 the area). Simulations were performed to measure the probability distribution for the number of surviving species under four conditions (1,000 each): high and low competitive asymmetry and high and low structural disorder. The structural scale was held fixed. We used a maximum likelihood estimator (MLE) to measure the ensemble average survival probability (*α*) under those four conditions. Without competitive asymmetry (red and green X’s), the number of surviving species was heavily weighted toward the maximum possible number (8). With high competitive asymmetry, the probability distributions spread across all possible numbers of species with relatively little distinction between ordered and disordered systems (cyan and purple X’s). In all instances, the corresponding MLE fits are shown as solid lines. These results support the hypothesis that, regardless of structural scale, smaller environments hinder long-term species coexistence. In all simulations *L* = 75, ⟨*P*⟩ = 0.25, and Δ*x/R =* 3.5.

**SI Figure 4.**
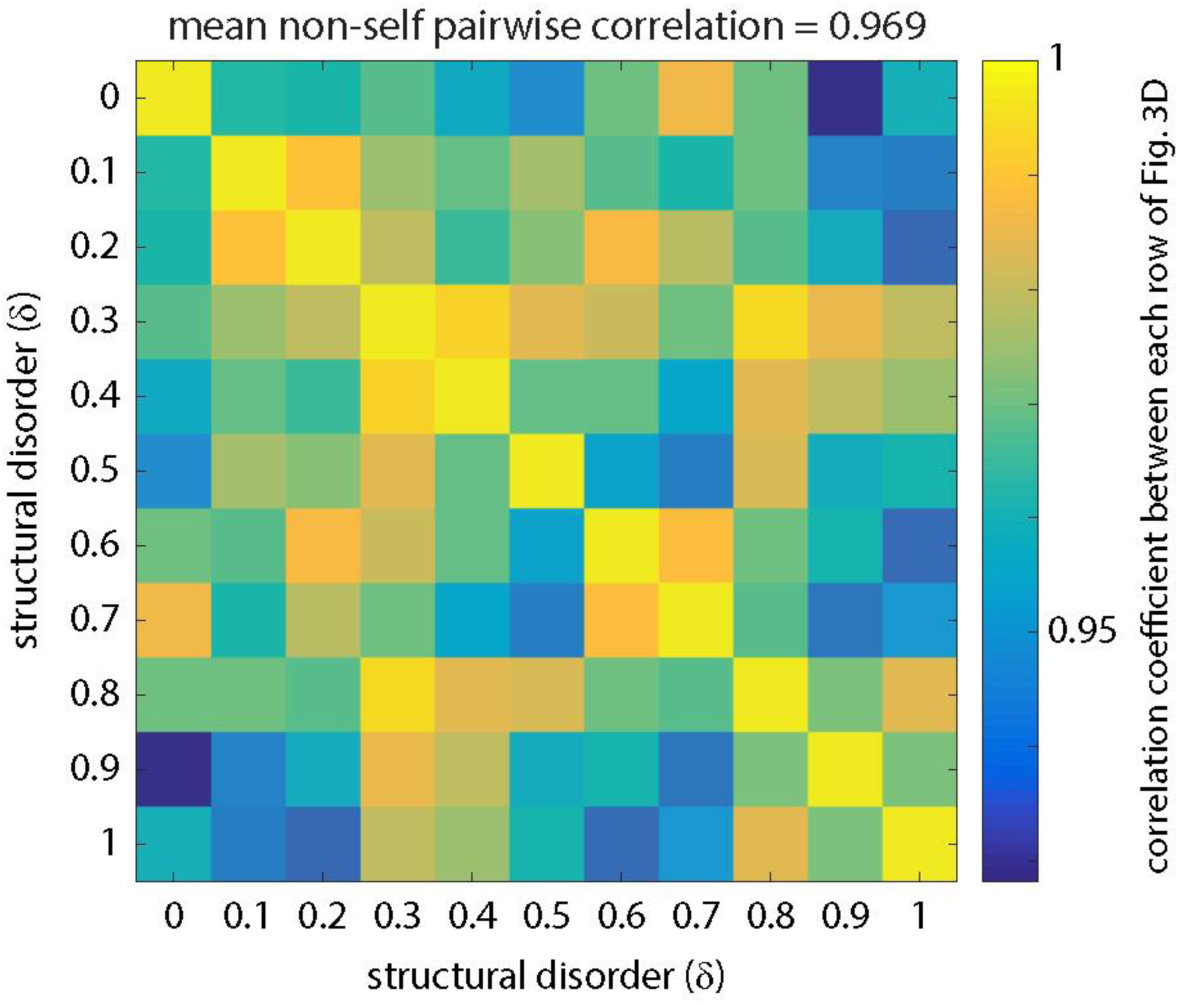
To determine the impact of steric disorder on the mean number of surviving species, independent of the effect of competitive asymmetry, we correlated each row of Fig. 3D with every other row of the same figure (55 unique correlations). All of those correlations were greater than 0.928, and the mean of all of those correlations was 0.969, meaning that over a range of steric disorder the relationship between mean number of surviving species and competitive asymmetry was quantitatively similar.

**SI Figure 5.**
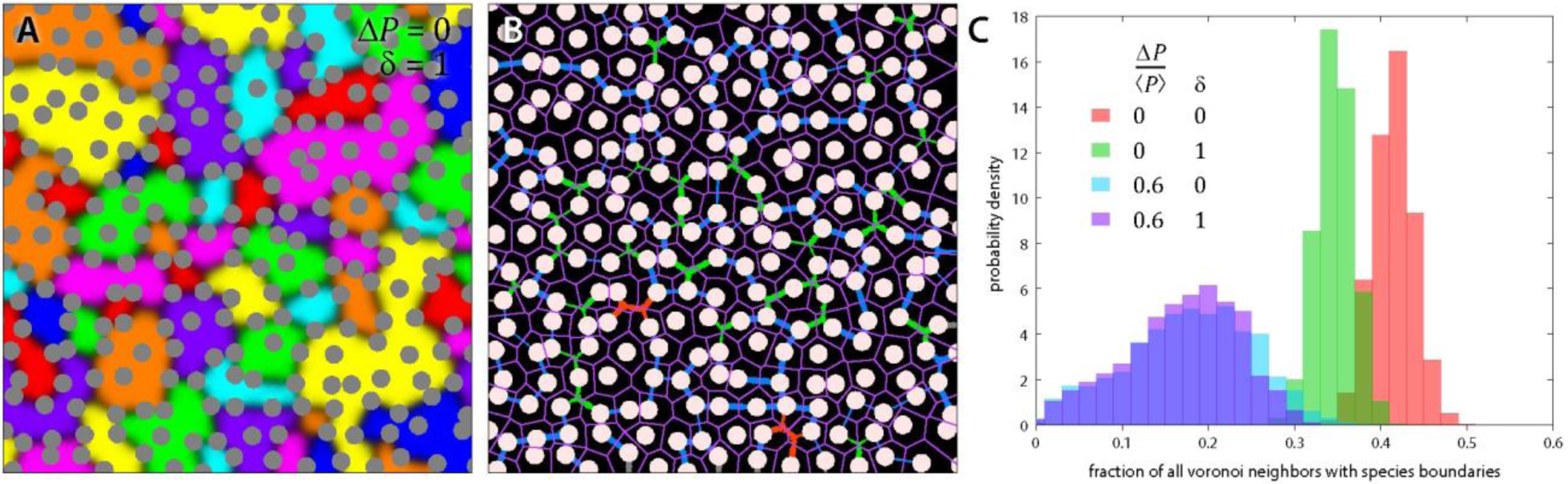
Using the same 4,000 simulations from Fig. 3, we examined the density of interspecies boundaries under those four conditions. **(A)** A sample image of 8 competing species in a disordered steric environment. Species establish spatial domains (solid colors) with dark boundaries between those domains where active competition takes places. **(B)** Across all simulations (though here for the same image as A), custom image analysis software examined the positions of each pillar (white) and the corresponding Voronoi tessellation (magenta tessellation) that indicates which pillars are Voronoi nearest neighbors. Image analysis algorithms segmented the interspecies boundaries between pillars and classified them according to how many pillars a boundary connected (blue = 2, green = 3, orange = 4). Approximately 2% of all connections were not Voronoi nearest neighbors (data not shown), and thus these connections were not used in this analysis. Boundaries that made contact with the edge of the simulation (gray) were not used in this analysis. **(C)** For each set of conditions, here shown as four colors (same colors as in Fig. 3B), we calculated the number of connections made by boundaries between Voronoi nearest neighbors as compared to the maximum possible number of boundaries (boundaries between all Voronoi nearest neighbors). While there is a notable difference between ordered and disordered connection density when competition is balanced (red and green), the salient difference is between balanced (red / green) and asymmetric (cyan / purple) competition. Systems with asymmetric competition have significantly fewer connections between objects, consistent with their higher abundance variability, and thus there is effectively less competition (i.e. fewer interfaces) in asymmetric systems.

**SI Figure 6.**
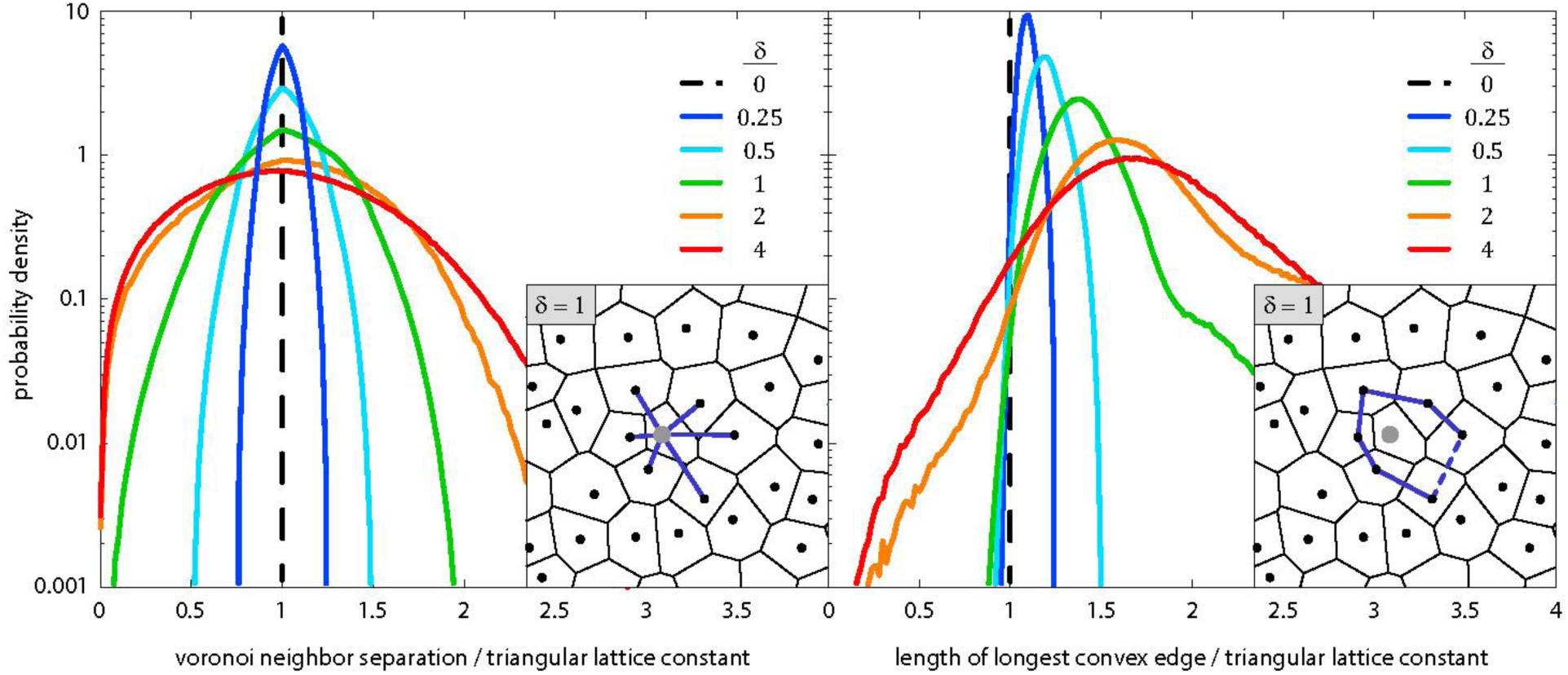
Local statistics of Voronoi tessellation as a function of the structural disorder. **(left)** Distributions of distances between Voronoi nearest neighbors. In a perfect triangular lattice this distribution is a delta-function (black dashed line). As disorder increases the distribution of nearest neighbor distances spreads out and a significant fraction of neighbors are found closer together than in a triangular lattice of the same overall density. The vast majority of interspecies boundaries are between Voronoi nearest neighbors. Thus widening of the distribution impacts whether a particular interspecies boundary is stable, because the maximum curvature and hence maximum competitive asymmetry that is stable at a boundary is set by the distance between the objects that the boundary connects. The inset shows an example of a Voronoi neighborhood and the local distances being measured (dark blue lines). **(right)** We examined hypothetical domains defined by the convex polygon around a steric object (see inset) – this defines a consistent region inside of which a competitor could stably exist (many other polygons could also be drawn). Across a large number of such polygon domains, we measured the distribution of longest edge lengths as a function of structural disorder. If the longest edge is less than the triangular lattice constant with the same overall density, then this domain is guaranteed to be more stable to competitive asymmetry than the corresponding ordered polygon. As disorder increases, the fraction of all such polygons that meet this more stringent stability condition increases, supporting the hypothesis that disorder should have a stabilizing effect when competition is asymmetric, contingent on there being a sufficiently large area and hence sufficient opportunities for such domains to exist.

**SI Figure 7.**
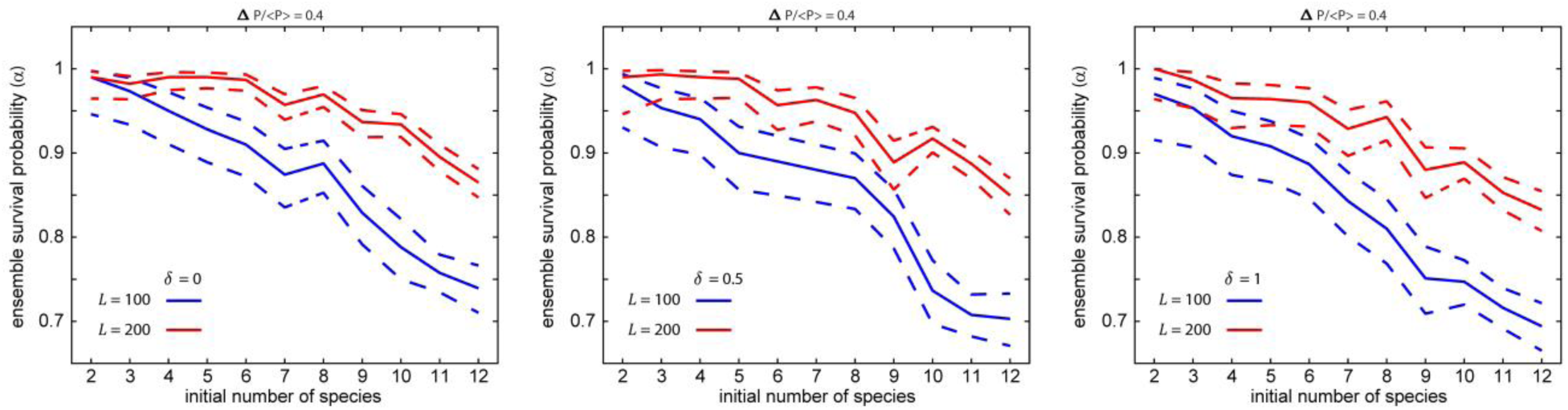
Same exact data as shown in Fig. 4B in the text, separated by value of *δ* and shown with 95% confidence intervals (dashed lines) determined by maximum-likelihood estimation. Variations caused by differences in *δ* are not significant, but variations caused by system-size differences are significant.

**SI Figure 8.**
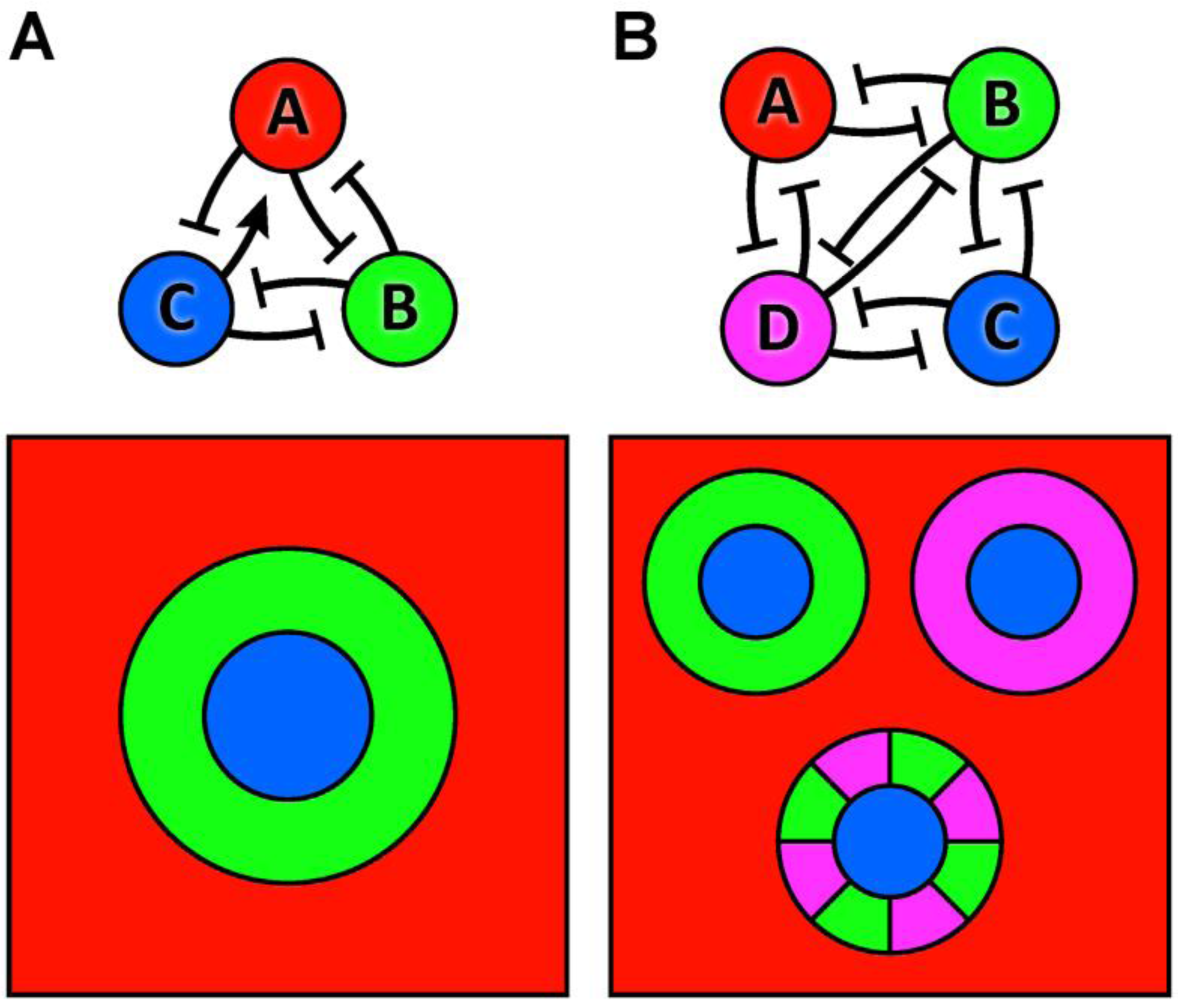
Spatial arrangements of species that support stability in non-all-to-all graphs. **(A)** The simplest example of a non-ATA graph (top) for which species abundances are stable in a structured environment – a single matrix element (arrow from *C* to *A*) breaks the ATA condition. The schematic (bottom) shows an arrangement of species that is stable, even though an interspecies boundary between *A* and *C* is not stable in any structured environment; this is the only arrangement that can stably support all three species for the graph shown. **(B)** A second, more complex example where four species can be arranged to yield stable abundances of all species, regardless of the interaction between species *A* and *C* (no connection shown). This graph admits three arrangements that allow *A* and *C* to coexist regardless of their interaction.

## REFERENCES

1. Daniel, R. The metagenomics of soil. Nat. Rev. Microbiol. 3, 470–478 (2005).

2. Stocker, R. & Seymour, J. R. Ecology and Physics of Bacterial Chemotaxis in the Ocean. Microbiol. Mol. Biol. Rev. 76, 792–812 (2012).

3. The Earth Microbiome Project Consortium et al. A communal catalogue reveals Earth’s multiscale microbial diversity. Nature 551, 457–463 (2017).

4. Melkonian, C. et al. Finding Functional Differences Between Species in a Microbial Community: Case Studies in Wine Fermentation and Kefir Culture. Front. Microbiol. 10, 1347 (2019).

5. Hutchinson, G. E. The Paradox of the Plankton. Am. Nat. 95, 137–145 (1961).

6. Gentry, A. H. Tree species richness of upper Amazonian forests. Proc. Natl. Acad. Sci. 85, 156–159 (1988).

7. Tilman, D. & Downing, J. A. Biodiversity and stability in grasslands. Nature 367, 363–365 (1994).

8. Ptacnik, R. et al. Diversity predicts stability and resource use efficiency in natural phytoplankton communities. Proc. Natl. Acad. Sci. 105, 5134–5138 (2008).

9. Blasius, B., Rudolf, L., Weithoff, G., Gaedke, U. & Fussmann, G. F. Long-term cyclic persistence in an experimental predator–prey system. Nature 577, 226–230 (2020).

10. Levi, T. et al. Tropical forests can maintain hyperdiversity because of enemies. Proc. Natl. Acad. Sci. 116, 581–586 (2019).

11. Kerr, B., Riley, M. A., Feldman, M. W. & Bohannan, B. J. M. Local dispersal promotes biodiversity in a real-life game of rock–paper–scissors. Nature 418, 171–174 (2002).

12. Hoek, T. A. et al. Resource Availability Modulates the Cooperative and Competitive Nature of a Microbial Cross-Feeding Mutualism. PLOS Biol. 14, e1002540 (2016).

13. Pennekamp, F. et al. Biodiversity increases and decreases ecosystem stability. Nature 563, 109–112 (2018).

14. Yu, X., Polz, M. F. & Alm, E. J. Interactions in self-assembled microbial communities saturate with diversity. ISME J. 13, 1602–1617 (2019).

15. Adler, P. B., HilleRisLambers, J. & Levine, J. M. A niche for neutrality. Ecol. Lett. 10, 95–104 (2007).

16. Tilman, D. Competition and Biodiversity in Spatially Structured Habitats. Ecology 75, 2–16 (1994).

17. Hardin, G. The Competitive Exclusion Principle. Science 131, 1292–1297 (1960).

18. Faust, K. et al. Microbial Co-occurrence Relationships in the Human Microbiome. PLoS Comput. Biol. 8, e1002606 (2012).

19. Tilman, D. Niche tradeoffs, neutrality, and community structure: A stochastic theory of resource competition, invasion, and community assembly. Proc. Natl. Acad. Sci. 101, 10854–10861 (2004).

20. Silvertown, J. Plant coexistence and the niche. Trends Ecol. Evol. 19, 605–611 (2004).

21. Godoy, O., Stouffer, D. B., Kraft, N. J. B. & Levine, J. M. Intransitivity is infrequent and fails to promote annual plant coexistence without pairwise niche differences. Ecology 98, 1193–1200 (2017).

22. Wiles, T. J. et al. Host Gut Motility Promotes Competitive Exclusion within a Model Intestinal Microbiota. PLOS Biol. 14, e1002517 (2016).

23. Hubbell, S. P. Neutral theory in community ecology and the hypothesis of functional equivalence. Funct. Ecol. 19, 166–172 (2005).

24. McGill, B. J. A test of the unified neutral theory of biodiversity. Nature 422, 881–885 (2003).

25. McGill, B. J., Maurer, B. A. & Weiser, M. D. EMPIRICAL EVALUATION OF NEUTRAL THEORY. Ecology 87, 1411–1423 (2006).

26. O’Dwyer, J. P. & Cornell, S. J. Cross-scale neutral ecology and the maintenance of biodiversity. Sci. Rep. 8, 10200 (2018).

27. Dickens, B., Fisher, C. K. & Mehta, P. Analytically tractable model for community ecology with many species. Phys. Rev. E 94, 022423 (2016).

28. Bunin, G. Ecological communities with Lotka-Volterra dynamics. Phys. Rev. E 95, 042414 (2017).

29. Obadia, B. et al. Probabilistic Invasion Underlies Natural Gut Microbiome Stability. Curr. Biol. 27, 1999-2006.e8 (2017).

30. Vega, N. M. & Gore, J. Stochastic assembly produces heterogeneous communities in the Caenorhabditis elegans intestine. PLOS Biol. 15, e2000633 (2017).

31. Hallett, L. M., Shoemaker, L. G., White, C. T. & Suding, K. N. Rainfall variability maintains grass-forb species coexistence. Ecol. Lett. 22, 1658–1667 (2019).

32. Allesina, S. & Levine, J. M. A competitive network theory of species diversity. Proc. Natl. Acad. Sci. 108, 5638–5642 (2011).

33. Niehaus, L. et al. Microbial coexistence through chemical-mediated interactions. Nat. Commun. 10, 2052 (2019).

34. Pande, S. et al. Fitness and stability of obligate cross-feeding interactions that emerge upon gene loss in bacteria. ISME J. 8, 953–962 (2014).

35. Goldford, J. E. et al. Emergent simplicity in microbial community assembly. Science 7 (2018).

36. Pacheco, A. R., Moel, M. & Segrè, D. Costless metabolic secretions as drivers of interspecies interactions in microbial ecosystems. Nat. Commun. 10, 103 (2019).

37. Butler, S. & O’Dwyer, J. P. Stability criteria for complex microbial communities. Nat. Commun. 9, 2970 (2018).

38. Posfai, A., Taillefumier, T. & Wingreen, N. S. Metabolic Trade-Offs Promote Diversity in a Model Ecosystem. Phys. Rev. Lett. 118, 028103 (2017).

39. Weiner, B. G., Posfai, A. & Wingreen, N. S. Spatial ecology of territorial populations. Proc. Natl. Acad. Sci. 116, 17874–17879 (2019).

40. Goyal, A. & Maslov, S. Diversity, Stability, and Reproducibility in Stochastically Assembled Microbial Ecosystems. Phys. Rev. Lett. 120, 158102 (2018).

41. Yurtsev, E. A., Conwill, A. & Gore, J. Oscillatory dynamics in a bacterial cross-protection mutualism. Proc. Natl. Acad. Sci. 113, 6236–6241 (2016).

42. Mougi, A. & Kondoh, M. Diversity of Interaction Types and Ecological Community Stability. Science 337, 349–351 (2012).

43. Levine, J. M., Bascompte, J., Adler, P. B. & Allesina, S. Beyond pairwise mechanisms of species coexistence in complex communities. Nature 546, 56–64 (2017).

44. Mickalide, H. & Kuehn, S. Higher-Order Interaction between Species Inhibits Bacterial Invasion of a Phototroph-Predator Microbial Community. Cell Syst. 9, 521-533.e10 (2019).

45. Friedman, J., Higgins, L. M. & Gore, J. Community structure follows simple assembly rules in microbial microcosms. Nat. Ecol. Evol. 1, 0109 (2017).

46. Kelsic, E. D., Zhao, J., Vetsigian, K. & Kishony, R. Counteraction of antibiotic production and degradation stabilizes microbial communities. Nature 521, 516–519 (2015).

47. Vetsigian, K., Jajoo, R. & Kishony, R. Structure and Evolution of Streptomyces Interaction Networks in Soil and In Silico. PLoS Biol. 9, e1001184 (2011).

48. Hart, S. P., Turcotte, M. M. & Levine, J. M. Effects of rapid evolution on species coexistence. Proc. Natl. Acad. Sci. 116, 2112–2117 (2019).

49. Liu, F. et al. Interaction variability shapes succession of synthetic microbial ecosystems. Nat. Commun. 11, 309 (2020).

50. Di Giacomo, R. et al. Deployable micro-traps to sequester motile bacteria. Sci. Rep. 7, 45897 (2017).

51. Großkopf, T. & Soyer, O. S. Synthetic microbial communities. Curr. Opin. Microbiol. 18, 72–77 (2014).

52. Hynes, W. F. et al. Bioprinting microbial communities to examine interspecies interactions in time and space. Biomed. Phys. Eng. Express 4, 055010 (2018).

53. Cremer, J. et al. Effect of flow and peristaltic mixing on bacterial growth in a gut-like channel. Proc. Natl. Acad. Sci. 113, 11414–11419 (2016).

54. Yawata, Y., Nguyen, J., Stocker, R. & Rusconi, R. Microfluidic Studies of Biofilm Formation in Dynamic Environments. J. Bacteriol. 198, 2589–2595 (2016).

55. Ley, R. E., Peterson, D. A. & Gordon, J. I. Ecological and Evolutionary Forces Shaping Microbial Diversity in the Human Intestine. Cell 124, 837–848 (2006).

56. Stein, R. R. et al. Ecological Modeling from Time-Series Inference: Insight into Dynamics and Stability of Intestinal Microbiota. PLoS Comput. Biol. 9, e1003388 (2013).

57. Tropini, C., Earle, K. A., Huang, K. C. & Sonnenburg, J. L. The Gut Microbiome: Connecting Spatial Organization to Function. Cell Host Microbe 21, 433–442 (2017).

58. Wardle, D. A. The influence of biotic interactions on soil biodiversity. Ecol. Lett. 9, 870–886 (2006).

59. Strayer, D. L., Power, M. E., Fagan, W. F., Pickett, S. T. A. & Belnap, J. A Classification of Ecological Boundaries. BioScience 53, 723 (2003).

60. McNally, L. et al. Killing by Type VI secretion drives genetic phase separation and correlates with increased cooperation. Nat. Commun. 8, 14371 (2017).

61. Xiong, L., Cooper, R. & Tsimring, L. S. Coexistence and Pattern Formation in Bacterial Mixtures with Contact-Dependent Killing. Biophys. J. 114, 1741–1750 (2018).

62. Russell, A. B., Peterson, S. B. & Mougous, J. D. Type VI secretion system effectors: Poisons with a purpose. Nat. Rev. Microbiol. 12, 137–148 (2014).

63. Drissi, F., Buffet, S., Raoult, D. & Merhej, V. Common occurrence of antibacterial agents in human intestinal microbiota. Front. Microbiol. 6, 1–8 (2015).

64. Cordero, O. X. et al. Ecological Populations of Bacteria Act as Socially Cohesive Units of Antibiotic Production and Resistance. Science 337, 1228–1231 (2012).

65. Czaran, T. L., Hoekstra, R. F. & Pagie, L. Chemical warfare between microbes promotes biodiversity. Proc. Natl. Acad. Sci. 99, 786–790 (2002).

66. Boyer, F., Fichant, G., Berthod, J., Vandenbrouck, Y. & Attree, I. Dissecting the bacterial type VI secretion system by a genome wide in silico analysis: what can be learned from available microbial genomic resources? BMC Genomics 10, 104 (2009).

67. Contento, L., Hilhorst, D. & Mimura, M. Ecological invasion in competition–diffusion systems when the exotic species is either very strong or very weak. J. Math. Biol. 77, 1383–1405 (2018).

68. Holmes, E. E., Lewis, M. A., Banks, J. E. & Veit, R. R. Partial Differential Equations in Ecology: Spatial Interactions and Population Dynamics. Ecology 75, 17–29 (1994).

69. Momeni, B., Xie, L. & Shou, W. Lotka-Volterra pairwise modeling fails to capture diverse pairwise microbial interactions. eLife 6, e25051 (2017).

70. Vallespir Lowery, N. & Ursell, T. Structured environments fundamentally alter dynamics and stability of ecological communities. Proc. Natl. Acad. Sci. 116, 379–388 (2019).

71. Reichenbach, T., Mobilia, M. & Frey, E. Mobility promotes and jeopardizes biodiversity in rock– paper–scissors games. Nature 448, 1046–1049 (2007).

72. Reichenbach, T., Mobilia, M. & Frey, E. Noise and Correlations in a Spatial Population Model with Cyclic Competition. Phys. Rev. Lett. 99, 238105 (2007).

73. Feng, S.-S. & Qiang, C.-C. Self-organization of five species in a cyclic competition game. Phys. Stat. Mech. Its Appl. 392, 4675–4682 (2013).

74. Park, J., Do, Y. & Jang, B. Multistability in the cyclic competition system. Chaos Interdiscip. J. Nonlinear Sci. 28, 113110 (2018).

75. Jiang, L.-L., Zhou, T., Perc, M. & Wang, B.-H. Effects of competition on pattern formation in the rock-paper-scissors game. Phys. Rev. E 84, 021912 (2011).

76. Reichenbach, T., Mobilia, M. & Frey, E. Coexistence versus extinction in the stochastic cyclic Lotka-Volterra model. Phys. Rev. E 74, 051907 (2006).

77. Reichenbach, T. & Frey, E. Instability of Spatial Patterns and Its Ambiguous Impact on Species Diversity. Phys. Rev. Lett. 101, 058102 (2008).

78. Avelino, P. P. et al. How directional mobility affects coexistence in rock-paper-scissors models. Phys. Rev. E 97, 032415 (2018).

79. Durrett, R. & Levin, S. Spatial Aspects of Interspecific Competition. Theor. Popul. Biol. 53, 30–43 (1998).

80. Okabe, A., Boots, B. & Sugihara, K. Spatial Tessellations: Concepts and Applications of Voronoi Diagrams. in Wiley Series in Probability and Mathematical Statistics (1992). doi: 10.2307/2687299.

81. Avelino, P. P., Menezes, J., de Oliveira, B. F. & Pereira, T. A. Expanding spatial domains and transient scaling regimes in populations with local cyclic competition. Phys. Rev. E 99, 052310 (2019).

82. Gallien, L., Zimmermann, N. E., Levine, J. M. & Adler, P. B. The effects of intransitive competition on coexistence. Ecol. Lett. 20, 791–800 (2017).

83. Sinervo, B. & Lively, C. M. The rock-paper-scissors game and the evolution of alternative male strategies. Nature 380, 240–243 (1996).

84. Kirkup, B. C. & Riley, M. A. Antibiotic-mediated antagonism leads to a bacterial game of rock-paper-scissors in vivo. Nature 428, 412–414 (2004).

85. Higgins, L. M., Friedman, J., Shen, H. & Gore, J. Co-occurring soil bacteria exhibit a robust competitive hierarchy and lack of non-transitive interactions. bioRxiv https://doi.org/10.1101/175737 (2017).

86. Kim, H. J., Boedicker, J. Q., Choi, J. W. & Ismagilov, R. F. Defined spatial structure stabilizes a synthetic multispecies bacterial community. Proc. Natl. Acad. Sci. 105, 18188–18193 (2008).

87. Coyte, K. Z., Schluter, J. & Foster, K. R. The ecology of the microbiome: networks, competition and stability. Science 350, 663–6 (2015).

88. Oliveira, N. M., Niehus, R. & Foster, K. R. Evolutionary limits to cooperation in microbial communities. Proc. Natl. Acad. Sci. 111, 17941–17946 (2014).

89. Huse, D. A. & Henley, C. L. Pinning and Roughening of Domain Walls in Ising Systems Due to Random Impurities. Phys. Rev. Lett. 54, 2708–2711 (1985).

90. Cammarota, C. & Biroli, G. Ideal Glass Transitions by Random Pinning. Proc. Natl. Acad. Sci. 109, 8850–8855 (2012).

91. Arumugam, S., Petrov, E. P. & Schwille, P. Cytoskeletal pinning controls phase separation in multicomponent lipid membranes. Biophys. J. 108, 1104–1113 (2015).

92. Gralka, M. & Hallatschek, O. Environmental heterogeneity can tip the population genetics of range expansions. eLife 8, e44359 (2019).

93. Fu, X., Cueto-Felgueroso, L. & Juanes, R. Thermodynamic coarsening arrested by viscous fingering in partially miscible binary mixtures. Phys. Rev. E 94, 1–5 (2016).

94. Martinez-Garcia, R., Nadell, C. D., Hartmann, R., Drescher, K. & Bonachela, J. A. Cell adhesion and fluid flow jointly initiate genotype spatial distribution in biofilms. PLoS Comput. Biol. 14, e1006094 (2018).

95. Siryaporn, A., Kim, M. K., Shen, Y., Stone, H. A. & Gitai, Z. Colonization, competition, and dispersal of pathogens in fluid flow networks. Curr. Biol. 25, 1201–1207 (2015).

96. Borer, B., Tecon, R. & Or, D. Spatial organization of bacterial populations in response to oxygen and carbon counter-gradients in pore networks. Nat. Commun. 9, (2018).

97. Cornforth, D. M. & Foster, K. R. Competition sensing: the social side of bacterial stress responses. Nat. Rev. Microbiol. 11, 285–293 (2013).

98. Mavridou, D. A. I., Gonzalez, D., Kim, W., West, S. A. & Foster, K. R. Bacteria Use Collective Behavior to Generate Diverse Combat Strategies. Curr. Biol. 28, 1–11 (2018).

99. Lowery, N. V., McNally, L., Ratcliff, W. C. & Brown, S. P. Division of labor, bet hedging, and the evolution of mixed biofilm investment strategies. mBio 8, (2017).

100. Cui, W., Marsland, R. & Mehta, P. Diverse communities behave like typical random ecosystems. http://biorxiv.org/lookup/doi/10.1101/596551 (2019) doi: 10.1101/596551.

